# Updating the steady state model of C_4_ photosynthesis

**DOI:** 10.1101/2021.03.13.435281

**Authors:** Susanne von Caemmerer

## Abstract

C_4_ plants play a key role in world agriculture. For example, C_4_ crops such as maize and sorghum are major contributors to both first and third world food production and the C_4_ grasses sugarcane; miscanthus and switchgrass are major plant sources of bioenergy. In the challenge to manipulate and enhance C_4_ photosynthesis, steady state models of leaf photosynthesis provide and important tool for gas exchange analysis and thought experiments that can explore photosynthetic pathway changes. Here the C_4_ photosynthetic model by von Caemmerer and Furbank (1999) has been updated with new kinetic parameterisation and temperature dependencies added. The parameterisation was derived from experiments on the C_4_ monocot, *Setaria viridis*, which for the first time provides a cohesive parametrisation. Mesophyll conductance and its temperature dependence have also been included, as this is an important step in the quantitative correlation between the initial slope of the CO_2_ response curve of CO_2_ assimilation and in vitro PEP carboxylase activity. Furthermore, the equations for chloroplast electron transport have been updated to include cyclic electron transport flow and equations have been added to calculate electron transport rate from measured CO_2_ assimilation rates.

**Highlight:** The C_4_ photosynthesis model by von Caemmerer and Furbank (1999) has been updated. It now includes temperature dependencies and equations to calculate electron transport rate from measured CO_2_ assimilation rates.

## Introduction

To meet the challenge of increasing crop yield for a growing world population, it has become apparent that photosynthetic efficiency and capacity must be increased per unit leaf area to improve yield potential (Long *et al*., 2015). High yields from C_4_ crops have stimulated considerable interest in the C_4_ photosynthetic pathway which is characterised by high photosynthetic rate and high nitrogen and water use efficiency relative to plants with the C_3_ photosynthetic pathway (Mitchell and Sheehy, 2006). In the challenge to increase photosynthetic rate per leaf area steady state models of leaf photosynthesis provide an important tool for gas exchange analysis and thought experiments that can explore photosynthetic pathway changes (Long *et al*., 2015; Price *et al*., 2011; von Caemmerer and Evans, 2010; von Caemmerer and Furbank, 2016; von Caemmerer *et al*., 2003). The mathematical simplicity of these leaf level models has facilitated incorporation into higher order canopy, crop and earth system models (Rogers *et al*., 2017; Wu *et al*., 2018; Wu *et al*., 2019; Yin and Struik, 2009).

C_4_ photosynthesis requires the coordinated functioning of mesophyll and bundle-sheath cells of leaves and is characterised by a CO_2_ concentrating mechanism which allows Rubisco, located in the bundle sheath cells, to operate at high CO_2_ partial pressures. This overcomes the low affinity Rubisco has for CO_2_ and largely inhibits its oxygenation reaction, reducing photorespiration rates. In the mesophyll, CO_2_ is initially fixed by phosphoenolpyruvate (PEP) carboxylase into C_4_ acids, which are then decarboxylated in the bundle sheath to supply CO_2_ for Rubisco. Both the structure of the bundle-sheath wall (which has a low permeability to CO_2_) and the relative biochemical capacities of the C_3_ cycle in the bundle sheath and C_4_ acid cycle (which operates across the mesophyll bundle-sheath interface) contribute to the high CO_2_ partial pressure in the bundle sheath. The biochemistry of the C_4_ photosynthetic pathway is not unique and three main biochemical subtypes are recognised on a basis of the predominant decarboxylating enzyme: NADP-ME (NADP-dependent malic enzyme), NAD-ME (NAD-dependent malic enzyme) and PEPCK (PEP carboxykinase) (Hatch, 1987).

The first models to capture the C_4_ photosynthetic biochemistry were designed by Berry and Farquhar (1978) and (Peisker, 1979). The Berry and Farquhar model did not provide analytical solutions but was able to predict high bundles sheath CO_2_ partial pressures and its dependence on bundle sheath conductance. Many of the gas exchange characteristics of C_4_ photosynthesis observed with intact leaves could be predicted by these models. Collatz *et al*. (1992) and von Caemmerer and Furbank (1999) have revised and expanded these original models with analytical solutions.

C_4_ models have not been used as frequently as the C_3_ models so less data relating leaf biochemistry with gas exchange is available in the literature. Massad *et al*. (2007) have parameterised the model by von Caemmerer (2000) for *Zea mays* and developed the first temperature dependencies for key parameters. Fitting routines have also been developed (Bellasio *et al*., 2016; Bellasio *et al*., 2017; Zhou *et al*., 2019).

Here an update of the C_4_ photosynthetic model by von Caemmerer and Furbank (1999) and von Caemmerer (2000) is provided with new parameterisation and temperature dependencies derived from experiments on the C_4_ monocot species *Setaria viridis* (green foxtail millet), a NADP-malic enzyme type which is closely related to agronomically important C_4_ crops. It has become a popular model species due to its rapid generation time, small stature, high seed production, diploid status and small sequenced and publicly available genome and it can be readily transformed (Alonso-Cantabrana *et al*., 2018; Brutnell *et al*., 2010; Doust, 2007; Ermakova *et al*., 2019; Li and Brutnell, 2011; Osborn *et al*., 2016).

### The basic model equations

Figure 1 shows a schematic representation of the proposed carbon fluxes in C_4_ photosynthesis. After diffusion of CO_2_ across the mesophyll cell interface CO_2_ is converted to HCO_3-_ by carbonic anhydrase, CA, which is fixed by PEP carboxylase, PEPC into C_4_ acids, which diffuse to and are decarboxylated in the bundle sheath. Rubisco and the complete C_3_ photosynthetic pathway are located in the bundle-sheath cells, bounded by a relatively gas tight cell wall such that the C_3_ cycle relies almost entirely on C_4_ acid decarboxylation as its source of CO_2_.

**Figure 1.**
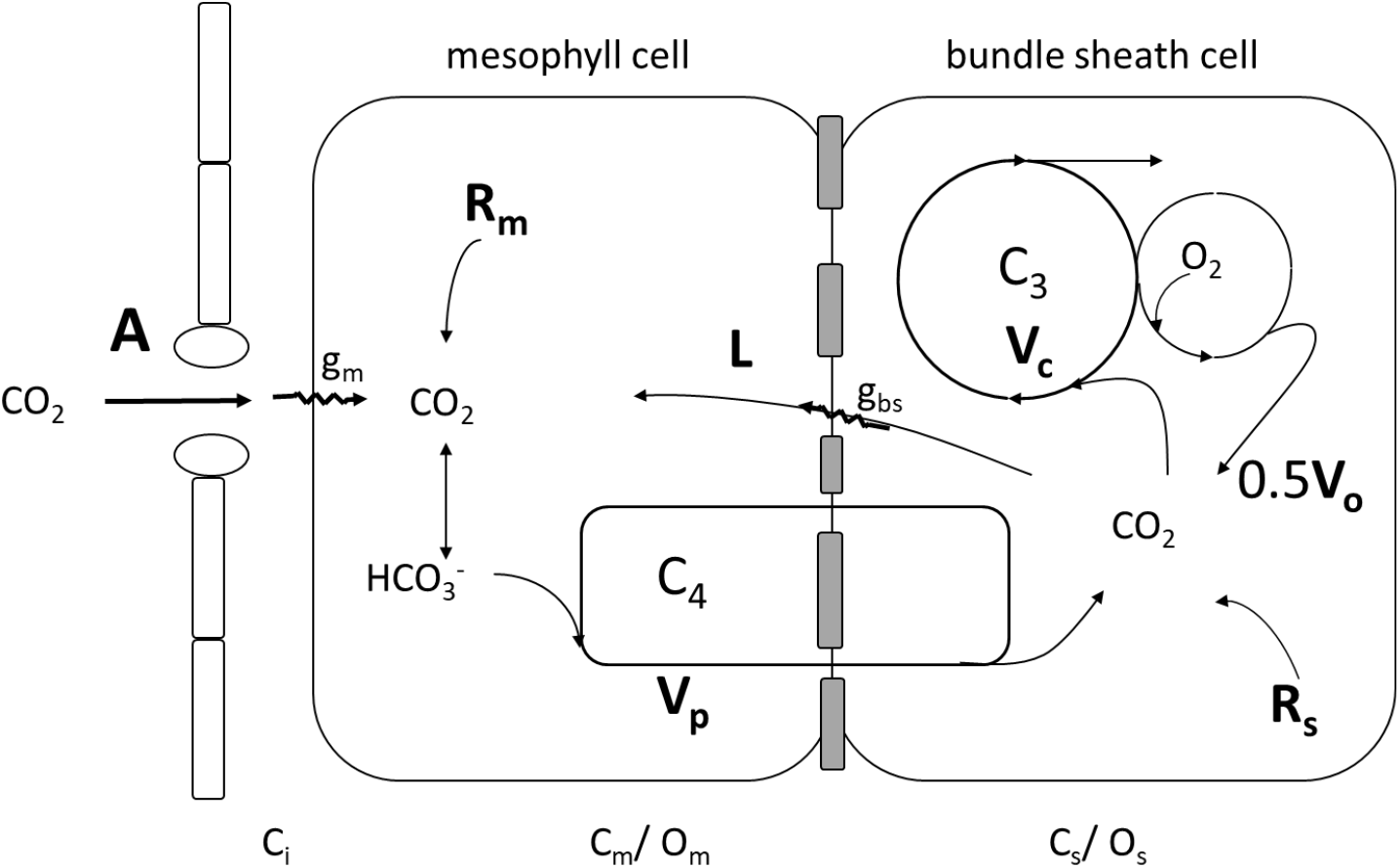
Schematic representing the main features of the C_4_ photosynthetic pathway. CO_2_ diffuses into the mesophyll where it is converted to HCO_3-_ and fixed by PEP carboxylase at the rate *V*_p_. In the steady state C_4_ acid decarboxylation occurs at the same rate. CO_2_ released in the bundle sheath either leaks out of the bundle sheath at (*L*) or is fixed by Rubisco (*V*_c_). In the photosynthetic carbon oxidation cycle CO_2_ is released at the half the oxygenation rate (*V*_o_). CO_2_ is also released by respiration (*R*_m_, *R*_s_) in mesophyll and bundle sheath respectively. Electron transport components are not shown.

The net rate of CO_2_ fixation for C_4_ photosynthesis can be given by two equations. The first describes Rubisco carboxylation in the bundle sheath. Since all carbon fixed into sugars ultimately must be fixed by Rubisco, overall CO_2_ assimilation, A, can be given by

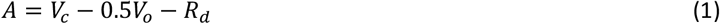

where *V*_c_ and *V*_o_ are the rates of Rubisco carboxylation and oxygenation and *R*_d_ is the rate of mitochondrial respiration not associated with photorespiration.

Mitochondrial respiration may occur in the mesophyll as well as in the bundle sheath. As Rubisco may more readily refix CO_2_ released in the bundle sheath, *R*_d_ is described by its mesophyll and bundle-sheath components

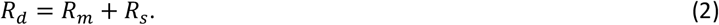

CO_2_ assimilation rate, *A*, can also be written in terms of the mesophyll reactions as

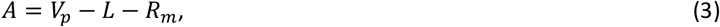

where *V*_*p*_ is the rate of PEP carboxylation, *R*_m_ is the mitochondrial respiration occurring in the mesophyll and *L* is the rate of CO_2_ leakage from the bundle sheath to the mesophyll (Figure 1). This assumes that in the steady state the rate of PEP carboxylation and the rate of C_4_ acid decarboxylation are equal.

The leak rate, *L*, is given by

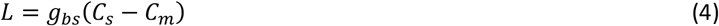

where *g*_*bs*_ is the conductance to CO_2_ leakage and is determined by the properties of the bundle-sheath cell wall; C_s_ and C_m_ are the bundle sheath and mesophyll CO_2_ partial pressures. It is assumed that there is a negligible amount of HCO_3-_ leakage from the bundle sheath since the HCO_3-_ pool should be small due to the absence of carbonic anhydrase activity in the cytosol of these cells (Farquhar, 1983; Jenkins *et al*., 1989; Ludwig *et al*., 1998).

The C_4_ cycle consumes additional energy during the regeneration of PEP and leakage of CO_2_ from the bundle sheath is an energy cost to the leaf. This represents a compromise between retaining CO_2_, allowing efflux of O_2_ and permitting metabolites to diffuse in and out at rates fast enough to support the rate of CO_2_ fixation (Hatch and Osmond, 1976; Raven, 1977). The CO_2_ leak rate depends upon the balance between the rates of PEP carboxylation and Rubisco activity and the conductance of the bundle sheath to CO_2_.

Leakiness (ϕ), a term coined by Farquhar (1983), defines leakage as a fraction of the rate of PEP carboxylation and thus describes the efficiency of the C_4_ cycle

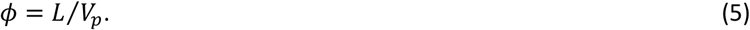

A related term “overcycling” has also been used (Furbank *et al*., 1990; Jenkins, 1989). Overcycling defines the leak rate as a fraction of CO_2_ assimilation rate and gives the fraction by which the flux through the C_4_ acid cycle has to exceed net CO_2_ assimilation rate

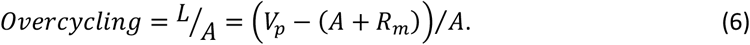

C_4_ photosynthesis can be either limited by the enzymatic rates of PEP carboxylase and Rubisco or by the irradiance and the capacity of chloroplast electron transport which supports the regeneration of PEP and RuBP.

### Enzyme limited rate equations

Many important features of the C_4_-model can be examined with the enzyme-limited rates, which are presumed to be appropriate under conditions of high irradiance. As is the case in C_3_ models of photosynthesis (Farquhar and von Caemmerer, 1982; Farquhar *et al*., 1980; von Caemmerer, 2000), Rubisco carboxylation at high irradiance can be described by its RuBP saturated rate

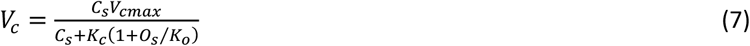

where O_s_ is the O_2_ partial pressure in the bundle sheath. Following the oxygenation of one mol of RuBP, 0.5 mol of CO_2_ is evolved in the photorespiratory pathway and the ratio of oxygenation to carboxylation can be expressed as

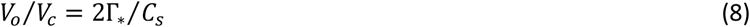

where *Γ*\* is the CO_2_ compensation point in a C_3_ plant in the absence of other mitochondrial respiration, and

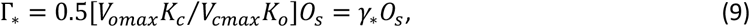

where the term in the bracket is the reciprocal of Rubisco specificity, *S*_c/o_ (Farquhar *et al*., 1980). In what follows the O_2_ dependence of *Γ*\* is highlighted since the O_2_ partial pressure in the bundle sheath may vary.

The Rubisco limited rate of CO_2_ assimilation can be derived from equations 1, 7, 8 and 9.

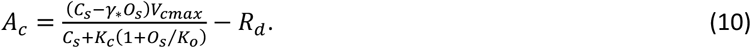

To derive an overall expression for CO_2_ assimilation rate as a function of mesophyll CO_2_ and O_2_ partial pressure, C_m_ and O_m_, one needs to derive an expression for C_s_ and O_s_. Equation 10 can be used to derive an expression for C_s_:

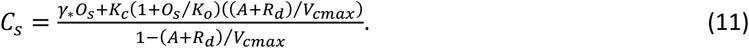

If *V*_cmax_ could be estimated accurately from biochemical measurements together with *A* it would provide a means of estimating bundle sheath CO_2_ partial pressure. One can also obtain an expression for C_s_ from equation 3 and 4:

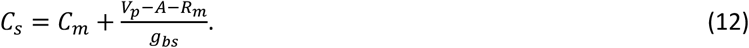

Photosystem II activity and O_2_ evolution in the bundle sheath varies widely amongst the C_4_ species. Some NADP-ME species such as *Zea mays* and *Sorghum bicolor* have little or none, whereas NADP-ME dicots and NAD and PCK species can have high PSII activity (Chapman *et al*., 1980; Hatch, 1987; Pfundel and Pfeffer, 1997). In S. viridis the amount of PSII activity depends on growth light environment (Ermakova *et al*., 2021). Because the bundle sheath is a fairly gas tight compartment this has implications for the steady state O_2_ partial pressure of the bundle sheath (Berry and Farquhar, 1978; Raven, 1977). Following Berry and Farquhar (1978), we assume that the net O_2_ evolution, *E*_o,_ in the bundle sheath cells equals its leakage, *L*_o_, out of the bundle sheath, that is

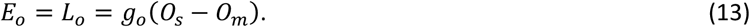

The conductance to leakage of O_2=_ across the bundle sheath, *g*_o_ can be related to the conductance to CO_2_by way of the ratio of diffusivities and solubilities by

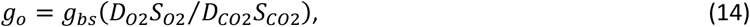

where *D*_*O*2_ and *D*_*CO*2_ are the diffusivities for O_2_ and CO_2_ in water, respectively and *S*_*O*2_ and are the respective Henry constants such that *S*_*CO*2_

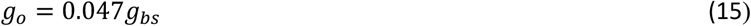

at 25° C (Berry and Farquhar, 1978; Farquhar, 1983). If *E*_o_=α*A*, where α (0<α>1) denotes the fraction of O_2_ evolution occurring in the bundle sheath, then *O*_s:_

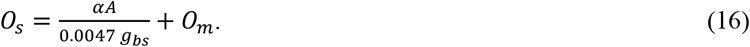

Like Berry and Farquhar (1978), it is assumed that a steady state balance exists between the rate of PEP carboxylation and the release of C_4_ acids in the bundle sheath. Furthermore, it is assumed that PEP carboxylation provides the rate limiting step and not, for example, the rate of hydration of CO_2_ by carbonic anhydrase. As PEP carboxylase utilises HCO_3_^−^ rather than CO_2_, hydration of CO_2_ is really the first step in carbon fixation in C_4_ species (Hatch and Burnell, 1990).

When CO_2_ is limiting the rate of PEP carboxylation is given by a Michaelis Menten equation

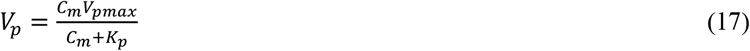

where *V*_pmax_ is the maximum PEP carboxylation rate, *K*_p_ is the Michaelis Menten constant for CO_2_. This assumes that the substrate PEP is saturating under these conditions. When the rate of PEP regeneration is limiting, for example by the capacity of pyruvate orthophosphate dikinase (PPDK) then

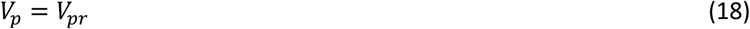

where *V*_pr_ is a constant (Peisker, 1986; Peisker and Henderson, 1992) and

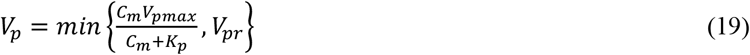

To obtain an overall rate equation for CO_2_ assimilation as a function of the mesophyll CO_2_ and O_2_partial pressures (*C*_m_ and *O*_m_) one combines equations (10), (12) and (16). The resulting expression is a quadratic of the form

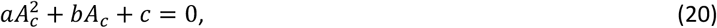

Where

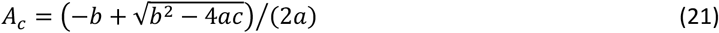

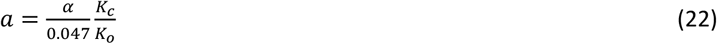

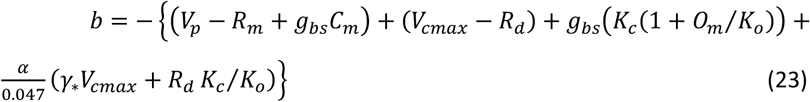

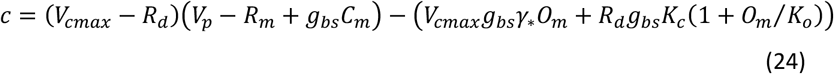

Equation (21) can be approximated by:

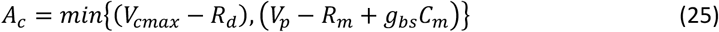

where min { } stands for minimum of.

At low CO_2_ partial pressures, CO_2_ assimilation rate can be approximated by

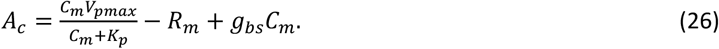

Under these conditions *A*_c_ is linearly related to the maximum PEP carboxylase activity, *V*_pmax_. The product *g*_bs_*C*_m_ is the inward diffusion of CO_2_ into the bundle sheath and because *g*_bs_ is low (0.003 mol m^−2^ s^−1^) the flux is only 0.3 μmol m^−2^ s^−1^ at C_m_ of a 100 μbar and can thus be ignored. At high CO_2_ partial pressures CO_2_ assimilation rate is given by either the maximal Rubisco activity, *V*_cmax_ or the rate of PEP regeneration (*V*_pr_).

### Light and electron transport limited rate equations

The energy requirements for the regeneration of RuBP in the bundle sheath are the same as in a C_3_ leaf (Farquhar *et al*., 1980; von Caemmerer, 2000). There is however the additional cost of 2 mol ATP for the regeneration of one mol of PEP from pyruvate in the mesophyll such that:

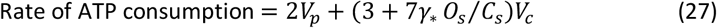

where (7*γ* O*_*s*_ / *C*_*s*_)*V*_*c*_ is the energy requirement due to photorespiration (since *V*_*o*_/*V*_*c*_ = *2*_*γ\**_ *O*_*s*_ /*C*_*s*_)(Berry and Farquhar, 1978). In the PCK type C_4_ species some of the ATP for PEP regeneration may come from the mitochondria such that the photosynthetic requirement may be less (for review see Furbank, 2011).

There is no net NADPH requirement by the C_4_ cycle itself, although for example in NADP-ME species NADPH consumed in the production of malate from OAA in the mesophyll is released in the bundle sheath during decarboxylation (Hatch and Osmond, 1976). This may have implications on the behaviour of C_4_ photosynthesis under fluctuating light environments (Krall and Pearcy, 1993; Kubásek *et al*., 2013). The rate of NADP consumption is given by the requirement of the C_3_ cycle:

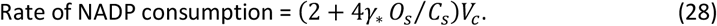

It is important to note that under most situations *C*_s_ is probably sufficiently large that the photorespiratory term in equations 27 and 28 can be ignored, but it does become relevant at low mesophyll CO_2_ partial pressures, or at very low light (Siebke *et al*., 1997).

NADPH and ATP are produced by chloroplast electron transport. The reduction of NADP^+^ to NADPH + H^+^ requires the transfer of 2 electrons through the whole chain electron transport which in turn requires 2 photons each at photosystem II (PSII) and Photosystem I (PSI). The generation of ATP can be coupled to the proton production via whole chain electron transport, or ATP can be generated via cyclic electron transport around PSI.

Photosystem II activity in the bundle sheath varies amongst C_4_ species with different C_4_ decarboxylation types. Presumably, when PSII is deficient or absent from the bundle sheath chloroplasts, some ATP is generated via cyclic photophosphorylation and 50% of the NADPH required for the reduction of PGA is derived from NADPH generated by NADP^+^ malic enzyme (Chapman *et al*., 1980). The remainder of the PGA must be exported to the mesophyll chloroplast where it is reduced and then returned to the bundle sheath (Hatch and Osmond, 1976). Measurements of metabolite pools of *Amaranthus edulis*, a NAD^+^ malic enzyme species, having PSII activity in the bundle sheath, suggest that it may also export a part of the PGA to the mesophyll for reduction (Leegood and von Caemmerer, 1988). It appears therefore that energy production and consumption is shared between mesophyll and bundle sheath cells more generally across decarboxylation types (von Caemmerer and Furbank, 2016).

A very simple approach was taken in the basic photosynthesis model. Electron transport is modelled as a whole, allocating a different fraction of it to the C_4_ and C_3_ cycle rather than compartmenting it to mesophyll and bundle sheath chloroplasts (von Caemmerer, 2000; von Caemmerer and Furbank, 1999).

That is whole chain linear electron transport

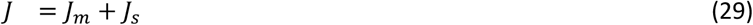

and *J*_*m*_ = *xJ* and *J*_*s*_ = (1 – *x*)*J* where 0<x>1. Because at most 2 out of 5 ATP are required in the mesophyll, *x* ≈ 0.4 (equation 27). This partitioning approach of electron transport has been adopted by subsequent users of the model (Kromdijk *et al*., 2010; Massad *et al*., 2007; Ubierna *et al*., 2011; Yin and Struik, 2009; Yin *et al*., 2011; Zhou *et al*., 2017). Peisker (1988) has modelled the optimisation of *x* at low light in some detail. See also Figure 4.22 in von Caemmerer (2000).

**Figure 4.**
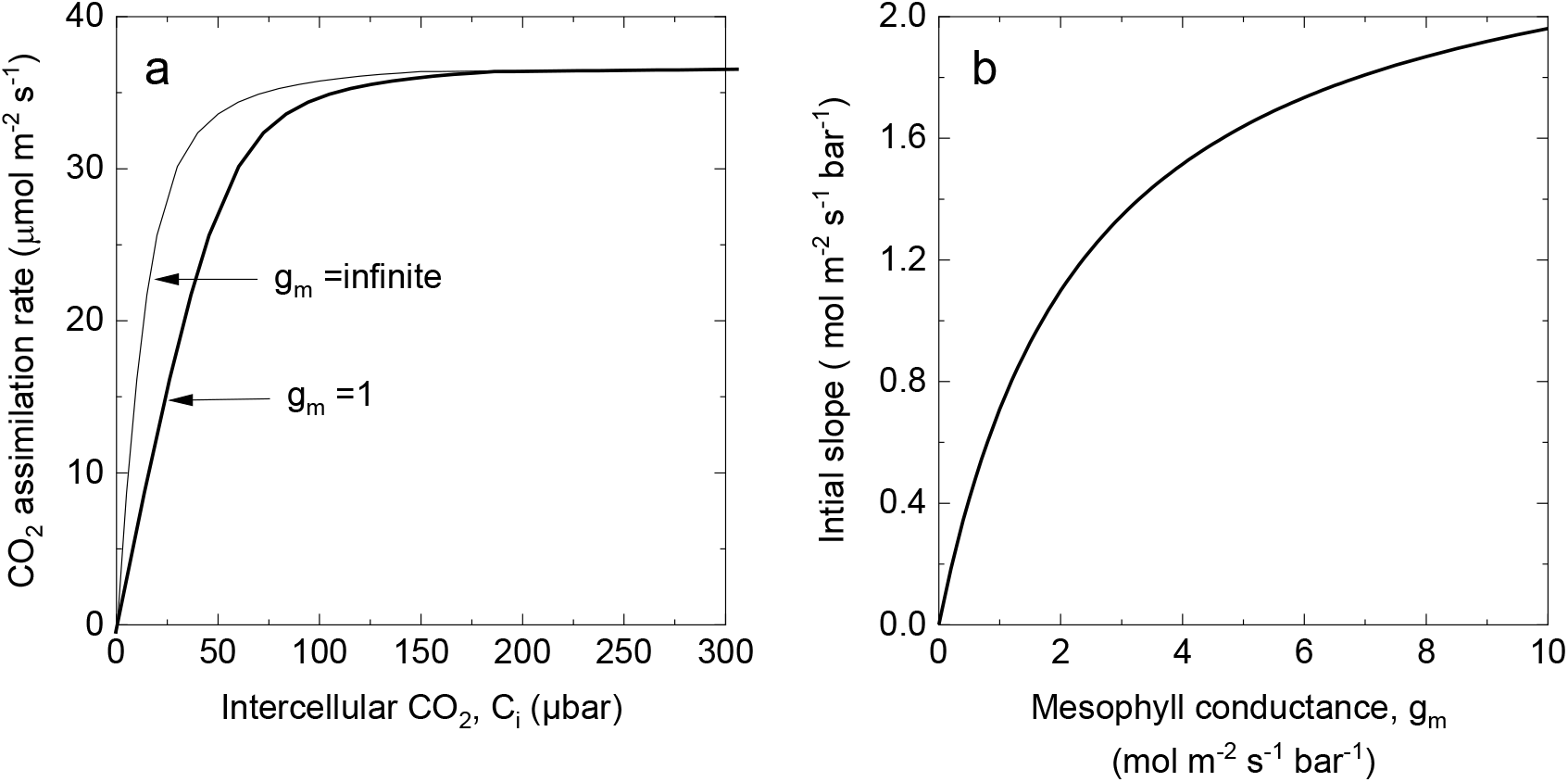
The effect of mesophyll conductance, g_m_ on the initial slope of the CO_2_ response curve. a) Modelled rate of CO_2_ assimilation as a function of intercellular CO_2_ partial pressure for the C_4_ photosynthetic pathway at 25 °C and an irradiance of 2000 µmol m^−2^ s^−1^modelled with g_m_ =1 mol m^−2^ s^−1^bar^−1^ or an infinite g_m._ b) Initial slope (equation x) as a function of mesophyll conductance. Model parameters are those given in Table 1.

New information exists for the calculation of the ATP requirement. Following Furbank et al. (1990), von Caemmerer and Furbank assumed a stoichiometry of 3 H^+^ per ATP produced and the operation of a Q-cycle (von Caemmerer, 2000; von Caemmerer and Furbank, 1999). Current models of rotational catalysis predict that the H^+^/ATP ratio is identical to the stoichiometric ratio of c-subunits to β-subunits which is c/β = 4.7 for spinach chloroplasts (Vollmar *et al*., 2009). However measured values are closer to 4 for the chloroplast enzyme (Petersen *et al*., 2012). If 4 H^+^ are required per ATP generated, it seems necessary to also have a functional Q-cycle which yields 3 H^+^ per linear electron flow. The proton production during cyclic electron flow is only 2 H^+^ per electron so that the overall proton production per electron is dependent on the balance of linear to cyclic electron flow (Yin and Struik, 2012). Following the derivations by Yin and Struik (2012), the rate of proton production from linear and cyclic electron flow is

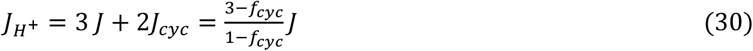

Since *J*_1_ = *J* + *J*_*cyc*_ = *J*/(1 – *f*_*cyc*_) where *J*_1_ is the electron flow out of photosystem 1 and f_cyc_ is the fraction of *J*_1_ that precedes via cyclic electron flow. For details see Figure 2. The rate of ATP production is given by

**Figure 2.**
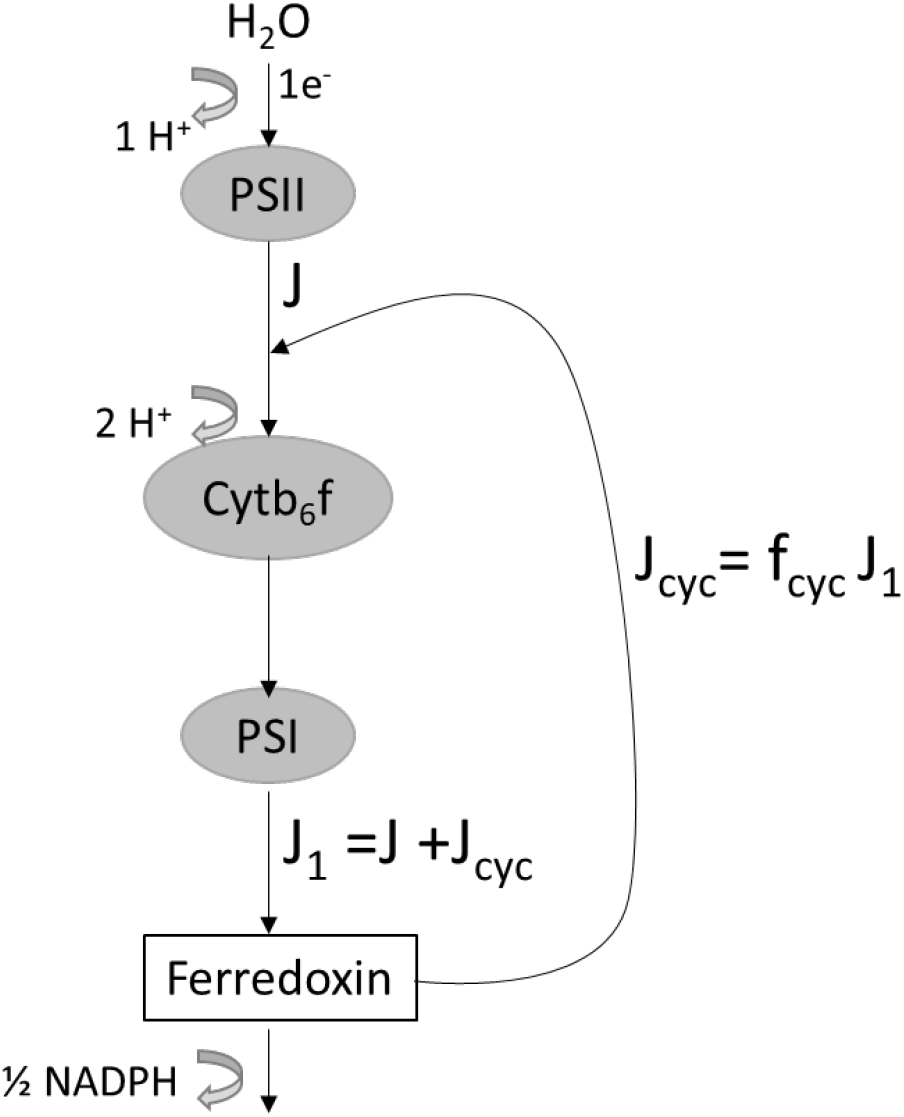
Scheme for linear and cyclic electron transport in the light reactions of photosynthesis. The arrows indictate electron transport. Thick curved arrows show the number of protons produced per electron transported. J denotes linear electron transport through photosystem II, J_1_ is electron transport trough photosystem I and J_cyc_ the rate of cyclic electron transport. The symbol f_cyc_ denotes the fraction of J_1_ that flows via the cyclic mode. The diagram has been adapted from Yin et al. (2004).

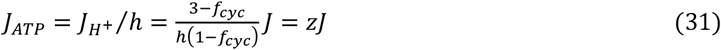

Where *h* is the number of protons required per ATP generated, which here is assumed to be 4 and *z* relates linear electron flow J to the rate of ATP production (Yin and Struik, 2012). It follows that

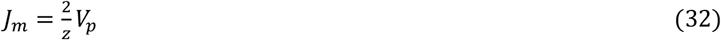

Where z=0.75 when there is no cyclic electron flow and z=1.25 when f_cyc_ =0.5.

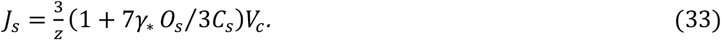

The relationship between the electron transport, J, and the absorbed irradiance that is used here is the same as that used previously where

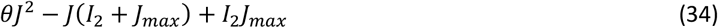

I_2_ is the photosynthetically useful light absorbed by PSII, J_max_ is the maximum electron transport and θ is an empirical curvature factor. 0.7 is a good average value for C_3_ species (Evans, 1989) but has not been explored for C_4_ species. *I*_2_ is related to incident irradiance by

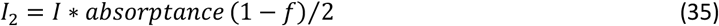

In sunlight the absorptance of leaves is commonly about 0.85 and *f* is to correct for spectral quality of the light (∼0.15 (Evans, 1987)). Ögren and Evans (1993) give a detailed discussion of the parameters of equation 16. The 2 in the denominator is because we assume half the light absorbed needs to reach each photosystem. This assumption may need to be considered as with an increase in cyclic electron flow commonly observed in C_4_ species less than half of the light may be absorbed by PSII.

Putting it all together gives a light and electron transport limited quadratic expression. From equations (3) and (1) one can derive two equations for an electron transport limited CO_2_ assimilation rate

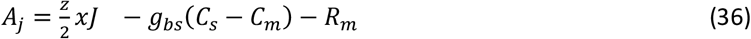

and

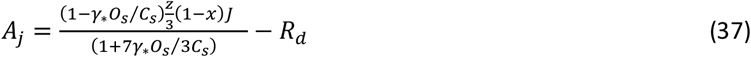

Equation (37) can be solved for the bundle sheath CO_2_ partial pressure and

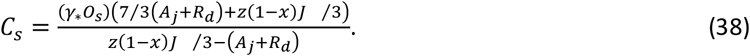

Combining equations (16), (36) and (36) then yields a quadratic expression of the form

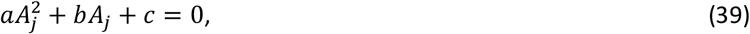

Where

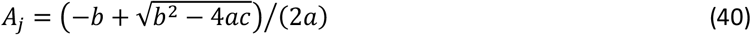

and

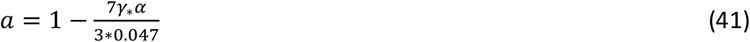

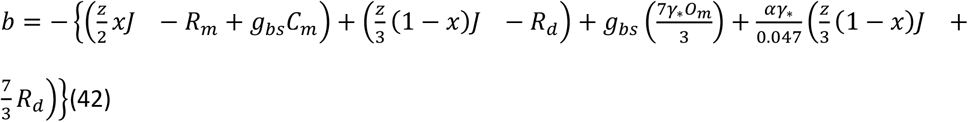

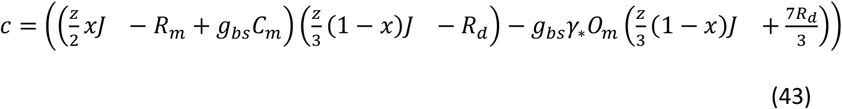

Equation (4.40) can be approximated by

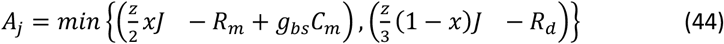

Where min { } stands for minimum of. Sometimes, when the equations are used to fit gas exchange measurements it is sufficient to use equation (44).

### Summary of equations

Equations (21) and (40) are the two basic equations of the C_4_ model and

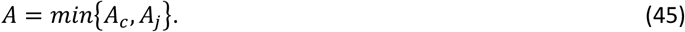

Peisker and Henderson (1992) pointed out that either the enzyme activity or the substrate regeneration rate can limit both Rubisco and PEP carboxylase reactions and that in theory four types of combinations of rate limitations are possible. In the way the electron transport limited equations are presented here, it is assumed that light or the electron transport capacity limit both PEP and RuBP regeneration rates simultaneously. In the model of C_3_ photosynthesis by Farquhar et al. (1980) and von Caemmerer and Farquhar (1981) it was assumed that the limitation of RuBP regeneration could be adequately modelled by an electron transport limitation without consideration of limitations by other PCR cycle enzymes. This is probably the case in most instances however in transgenic studies care needs to be taken. Transgenic tobacco with reduced sedoheptulose 1,7 bisposphatase (SBPase) regeneration of RuBP has been shown to be the more limiting step (Harrison *et al*., 2001; Harrison *et al*., 1998). In the case of C_4_ photosynthesis the possibility that PEP regeneration may also be limited by the enzyme activity of enzymes such as PPDK and malic enzyme at high irradiance has also been found in transgenic studies with *Flaveria bidentis* a C_4_ dicot (Furbank *et al*., 2001; Pengelly *et al*., 2012; Trevanion *et al*., 1997). C_4_ transgenic plants with altered RuPB regeneration capacity have not yet been reported on.

### An update on the parameterisation of the C_4_ photosynthesis model

This model is built on the same principal as the Farquhar et al. (1980) model and many of the model’s parameters can be assigned a priori and this is indicated in Table 1, leaving only key variables like V_cmax_ V_pmax_, V_pr_, and J_max_ to be assigned. It is however important to note that the kinetic constants of Rubisco from C_4_ species differ from those of C_3_ species and vary amongst the different C_4_ decarboxylation types (Badger *et al*., 1974; Ghannoum *et al*., 2005; Jordan and Ogren, 1981, 1983; Sharwood *et al*., 2016).

**Table 1.**
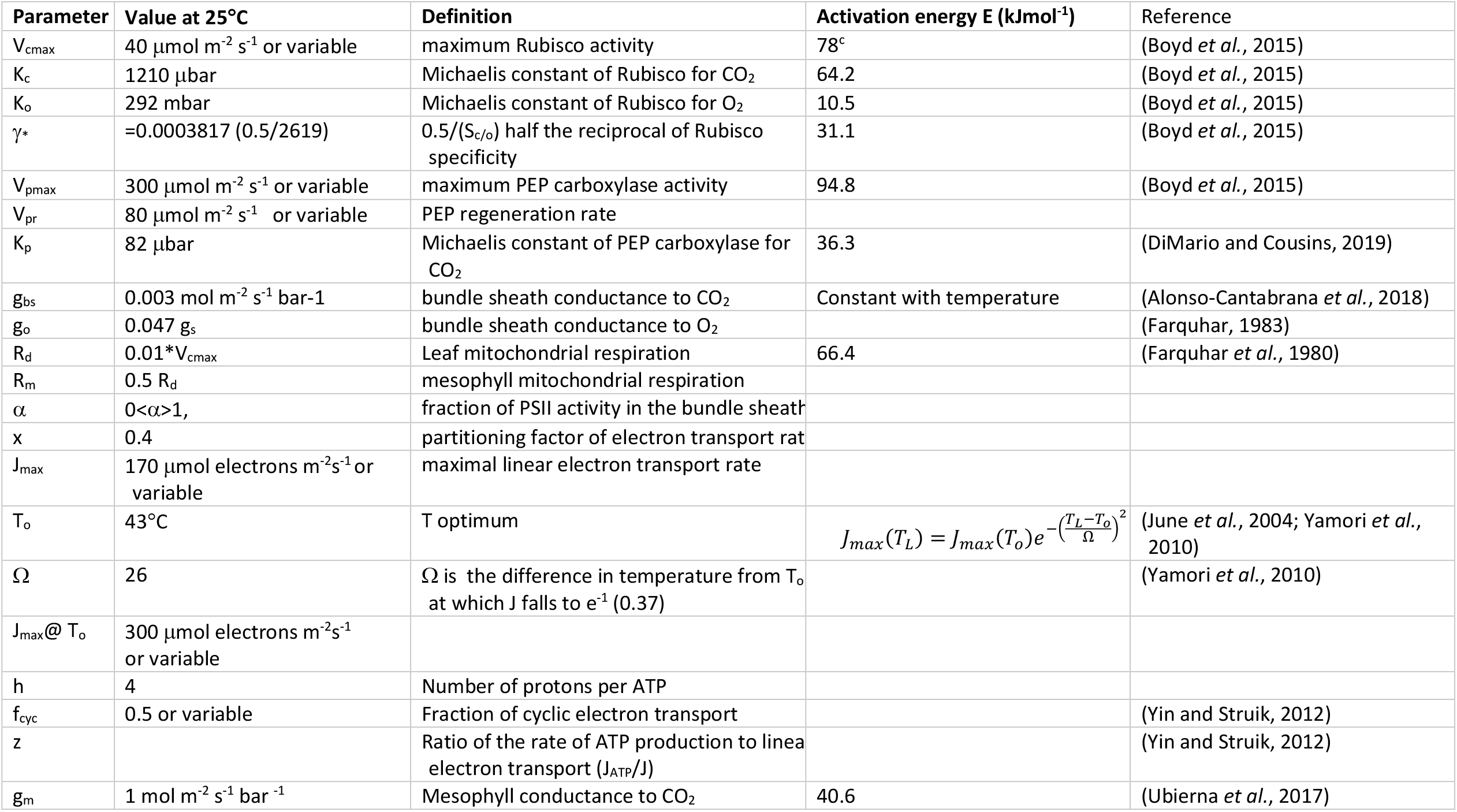
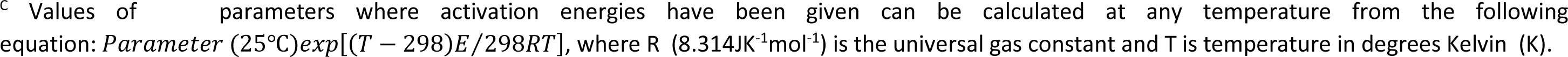
Photosynthetic parameters used in the model. When available values for *Setaria viridis* have been chosen.

| Parameter | Value at 25°C | Definition | Activation energy E (kJmol<sup>-1</sup>) | Reference |
| --- | --- | --- | --- | --- |
| V<sub>cmax</sub> | 40 µmol m<sup>-2</sup> s<sup>-1</sup> or variable | maximum Rubisco activity | 78<sup>c</sup> | (Boyd *et al.*, 2015) |
| K<sub>c</sub> | 1210 µbar | Michaelis constant of Rubisco for CO<sub>2</sub> | 64.2 | (Boyd *et al.*, 2015) |
| K<sub>o</sub> | 292 mbar | Michaelis constant of Rubisco for O<sub>2</sub> | 10.5 | (Boyd *et al.*, 2015) |
| γ* | =0.0003817 (0.5/2619) | 0.5/(S<sub>c/o</sub>) half the reciprocal of Rubisco specificity | 31.1 | (Boyd *et al.*, 2015) |
| V<sub>pmax</sub> | 300 µmol m<sup>-2</sup> s<sup>-1</sup> or variable | maximum PEP carboxylase activity | 94.8 | (Boyd *et al.*, 2015) |
| V<sub>pr</sub> | 80 µmol m<sup>-2</sup> s<sup>-1</sup> or variable | PEP regeneration rate |  |  |
| K<sub>p</sub> | 82 µbar | Michaelis constant of PEP carboxylase for CO<sub>2</sub> | 36.3 | (DiMario and Cousins, 2019) |
| g<sub>bs</sub> | 0.003 mol m<sup>-2</sup> s<sup>-1</sup> bar<sup>-1</sup> | bundle sheath conductance to CO<sub>2</sub> | Constant with temperature | (Alonso-Cantabrana *et al.*, 2018) |
| g<sub>o</sub> | 0.047 g<sub>s</sub> | bundle sheath conductance to O<sub>2</sub> |  | (Farquhar, 1983) |
| R<sub>d</sub> | 0.01*V<sub>cmax</sub> | Leaf mitochondrial respiration | 66.4 | (Farquhar *et al.*, 1980) |
| R<sub>m</sub> | 0.5 R<sub>d</sub> | mesophyll mitochondrial respiration |  |  |
| α | 0<α>1, | fraction of PSII activity in the bundle sheath |  |  |
| x | 0.4 | partitioning factor of electron transport rate |  |  |
| J<sub>max</sub> | 170 µmol electrons m<sup>-2</sup>s<sup>-1</sup> or variable | maximal linear electron transport rate |  |  |
| T<sub>o</sub> | 43°C | T optimum | $J_{max}(T_L) = J_{max}(T_o)e^{-\left(\frac{T_L-T_o}{\Omega}\right)^2}$ | (June *et al.*, 2004; Yamori *et al.*, 2010) |
| Ω | 26 | Ω is the difference in temperature from T<sub>o</sub> at which J falls to e<sup>-1</sup> (0.37) |  | (Yamori *et al.*, 2010) |
| J<sub>max@ T<sub>o</sub></sub> | 300 µmol electrons m<sup>-2</sup>s<sup>-1</sup> or variable |  |  |  |
| h | 4 | Number of protons per ATP |  |  |
| f<sub>cyc</sub> | 0.5 or variable | Fraction of cyclic electron transport |  | (Yin and Struik, 2012) |
| z |  | Ratio of the rate of ATP production to linear electron transport (J<sub>ATP</sub>/J) |  | (Yin and Struik, 2012) |
| g<sub>m</sub> | 1 mol m<sup>-2</sup> s<sup>-1</sup> bar<sup>-1</sup> | Mesophyll conductance to CO<sub>2</sub> | 40.6 | (Ubierna *et al.*, 2017) |

<sup>c</sup> Values of parameters where activation energies have been given can be calculated at any temperature from the following equation: $Parameter(25^{\circ}\text{C}) \exp[(T - 298)E/298RT]$ , where R ( $8.314\text{JK}^{-1}\text{mol}^{-1}$ ) is the universal gas constant and T is temperature in degrees Kelvin (K).

The C_4_ monocot species *Setaria viridis* (green foxtail millet), a NADP-malic enzyme type which is closely related to agronomically important C_4_ crops including *Setaria italica* (foxtail millet), *Z. mays* (maize), *Sorghum bicolor* (sorghum) and *Saccharum officinarum* (sugarcane) has been suggested as a new model species (Brutnell *et al*., 2010). It has become a popular model species due to its rapid generation time, small stature, high seed production, diploid status and small sequenced and publicly available genome and it can be readily transformed (Alonso-Cantabrana *et al*., 2018; Brutnell *et al*., 2010; Doust, 2007; Li and Brutnell, 2011; Osborn *et al*., 2016). This has led to excellent biochemical characterisation of *S. viridis* PEPC and Rubisco (Boyd *et al*., 2015; DiMario and Cousins, 2019). Most parameters were taken from Boyd et al. (2015), but the Michaelis Menten constant for CO_2_, *K*_p_, was updated with more recent measurements by Di Mario and Cousins (2019). It is important to note that PEPc fixes bicarbonate rather than CO_2_ and *K*_p_ is converted from measured values of K_m_ for HCO_3-_^1^. Here a cytosolic pH of 7.2 and pKa =6.12 was assumed (Hatch and Burnell, 1990). The precise value of cytosolic pH is unknown and if a pH of 7.4 is assumed *K*_p_ decreases from 82 µbar to 50 µbar. PEPc S. viridis RNAi line has been used to characterise bundle-sheath conductance to CO_2_ diffusion (*g*_*bs*_) and its temperature dependence (Alonso-Cantabrana *et al*., 2018). This has for the first time provided a cohesive parameter set and the associated temperature functions (Table 1).

It is best to estimate the temperature response of J_max_ from a series of light response curves made at high CO_2_ and different temperatures as was done by Massad et al (2007) for *Zea mays*. They used an Arrhenius function for their parameterisation. The simpler temperature function suggested by June et al. (2004) is used here to parametrise the temperature dependence of electron transport (Table 1) and the parameterisation for tobacco has been used (Yamori *et al*., 2010) since these experiments still need to be done for *S. viridis*. Sonawane et al. (2017), who characterised the temperature response of CO_2_ assimilation rate in a number of C_4_ grasses used this function to fit the saturated rate of CO_2_ assimilation measured at high irradiance. The tobacco values chosen here fit within the range values reported for these C_4_ grasses (Sonawane *et al*., 2017)

### Calculating the electron transport required to sustain CO_2_ assimilation

Von Caemmerer and Farquhar (1981) suggested that measurements of CO_2_ assimilation rate can be used to calculate the electron transport rate needed to support the CO_2_ assimilation rate. Equation 37 can be used in the same way. Using equation 39 to solve for J_t_ this results in the following quadratic equation:

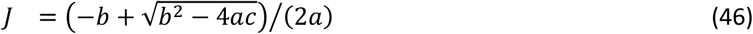

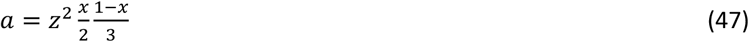

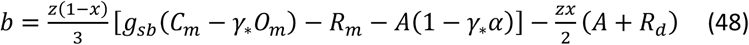

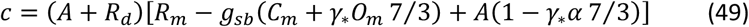

These equations were introduced by Ubierna et al.(2013) for linear electron flow only (z=0.75) and the assumption of 3 H^+^=/ATP.

## Model evaluation

### Modelled CO2 response of CO_2_ assimilation

In C_3_ species CO_2_ response curves are widely used to assess photosynthetic capacity (Ainsworth and Rogers, 2007; Sharkey *et al*., 2007; von Caemmerer, 2000; von Caemmerer and Farquhar, 1981). Figure 3 compares the model output of the Farquhar, von Caemmerer and Berry model of C_3_ photosynthesis (Farquhar *et al*., 1980) with the current C_4_ model presented here. In the C_3_ model the enzyme limited rate is dominated by Rubisco and its kinetic parameters at low CO_2_ and the electron transport capacity limits at high CO_2_ (Fig. 3a). In the C_4_ model it is also possible to distinguish an enzyme limited CO_2_ assimilation rate at high light (equations 20-25) and an electron transport limited rate (equations 37-44). However, the enzyme limited rate is determined by PEPC at low CO_2_ and Rubisco at high CO_2_. The electron transport limited rate can also determine the CO_2_ assimilation rate at high CO_2_ (Fig. 3b). Thus, it is more difficult to identify biochemical limitations to C_4_ photosynthesis.

**Figure 3.**
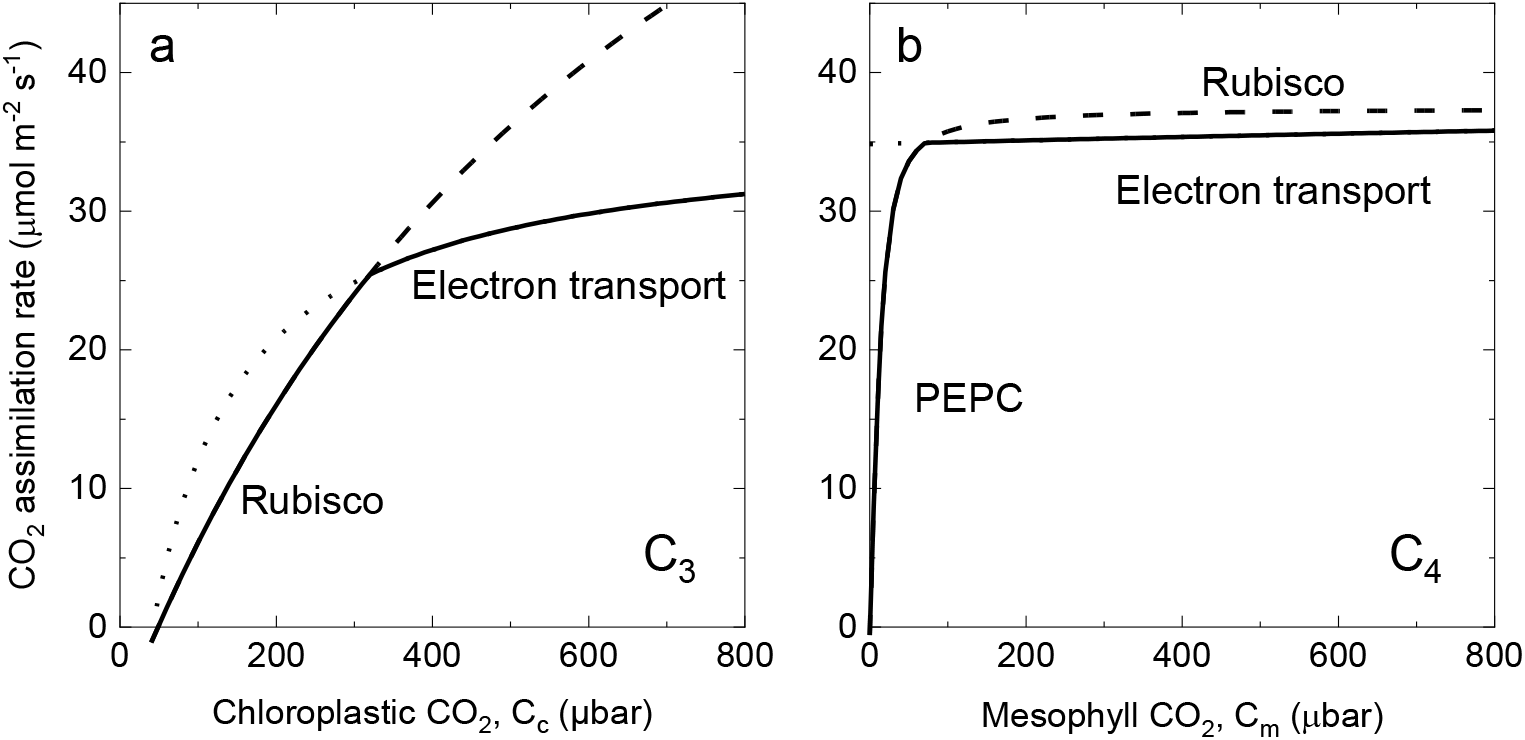
A comparison of modelled rates of CO_2_ assimilation rate as functions of partial pressures of CO_2_ for C_3_ (a) and C_4_ photosynthesis (b). c) Modelled rate of CO_2_ assimilation as a function of chloroplast CO_2_ partial pressure for the C_3_ photosynthetic pathway at 25 °C. The Rubisco limited (RuBP saturated rate) of CO_2_ assimilation has a dashed line extension at high CO_2_. The electron transport (RuBP regeneration) limited rate of CO_2_ assimilation has a dotted line extension at low CO_2_. The solid curve represents the minimum rate that is the actual rate of CO_2_ assimilation. A possible triose phosphate limitation at high CO_2_ is not shown. (Redrawn from von Caemmerer 2000). d) Modelled rate of CO_2_ assimilation as a function of mesophyll cytosolic CO_2_ partial pressure for the C_4_ photosynthetic pathway at 25 °C and an irradiance of 1500 µmol m^−2^ s^−1^. The enzyme limited CO_2_ assimilation rate (PEPC limitation at low CO_2_ and Rubisco limitation at high CO_2_ shows the Rubisco limited rate as a dashes line extension. The electron transport (RuBP and PEP regeneration) limited rate of CO_2_ assimilation has a dotted line extension at low CO_2_. The solid curve represents the minimum rate that is the actual rate of CO_2_ assimilation. Parameters used are given in Table 1.

Usually good correlations are found between in vitro Rubisco activity and the CO_2_ saturated rate of CO_2_ assimilation rate at high CO_2_ (Sonawane *et al*., 2017; Usuda, 1984; Usuda *et al*., 1984). The relationship should be almost one to one as Rubisco operates close to its saturated rate in vivo. In the study of *Flaveria bidentis* transgenics with varying reductions in Rubisco content show a slight curylinear relationship between Rubisco content and CO_2_ assimilation rate hinting at a possible electron transport limitation in wild type plants (Furbank *et al*., 1996; von Caemmerer *et al*., 1997). These studies have also provided evidence that Rubisco limits CO_2_ assimilation at high CO_2_. Transgenic plants with reduced Rubisco content showing a clear decline in CO_2_ assimilation rate at high CO_2_ (Pengelly *et al*., 2012; von Caemmerer *et al*., 1997). Recent photosynthetic engineering that increased Rubisco content in maize leaves resulted in an increase in CO_2_ saturated CO_2_ assimilation rate (Salesse-Smith *et al*., 2018).

Here the model has been tuned in such a way that at 25 °C electron transport rate is limiting CO_2_ assimilation at high CO_2_ and high irradiance. This balance can of course vary with growth conditions or species, but there is no straight forward technique to determine the limitation. Furthermore, the assumption has been made the electron transport capacity and PEP and RuBP regeneration generally co-limit. In transgenic studies where regeneration of the C_4_ cycle has been curtailed by molecular manipulation, it is clear that this also limits CO_2_ assimilation at high CO_2_ (Pengelly *et al*., 2012; Trevanion *et al*., 1997). In a study, transgenic *S. viridis* with overexpression of the Rieske iron sulphur protein in the cytochrome b_6_f complex had increased CO_2_ assimilation rates at ambient and high CO_2_ confirming that electron transport capacity can limit CO_2_ assimilation rate (Ermakova *et al*., 2019). The fact that electron transport rate limits CO_2_ assimilation rate at high CO_2_ means that a reduction in irradiance is also predicted to primarily affects the CO_2_ saturated rate of CO_2_ assimilation rather than the initial slope of the CO_2_ response curve except at low irradiance (Leegood and von Caemmerer, 1989; Pfeffer and Peisker, 1998).

There are three possible limitations to the initial slope of the CO_2_ response curve; the mesophyll conductance to CO_2_ diffusion from intercellular airspace to the mesophyll cytosol *g*_m_; the rate of CO_2_ hydration by carbonic anhydrase, CA, and the rate of PEP carboxylation. It is thought that most C_4_ leaves have sufficient CA for it not to be rate limiting (Cousins *et al*., 2008; Hatch and Burnell, 1990). However, studies with transgenic or mutant plants in *Flaveria bidenti*s, *Zea mays* and *Setaria viridis* have shown that when CA activity is greatly reduced a reduction in initial slope of the CO_2_ response is observed (Osborn *et al*., 2016; Studer *et al*., 2014; von Caemmerer *et al*., 2004).

The initial C_4_ photosynthesis models did not consider a diffusion limitation between the intercellular airspace and the mesophyll cytosol. In C_4_ species mesophyll conductance, g_m_, is likely to be proportional to mesophyll surface area exposed to intercellular airspace (Evans and von Caemmerer, 1996). The standard techniques used to quantify mesophyll conductance in C_3_ species such as combined measurements of gas exchange and chlorophyll fluorescence, or measurements of ^13^C isotope discrimination cannot be used in C_4_ species, however a new technique has been developed to measure mesophyll conductance in C_4_ species using C^18^O^16^O isotope discrimination (Barbour *et al*., 2016; Gillon and Yakir, 2000; Ogée *et al*., 2018; Osborn *et al*., 2016) and here the temperature dependence of g_m_ measured for *Setaria viridis* has been used for parameterisation of the model (Table 1, Ubierna *et al*., 2017).

The drop in CO_2_ partial pressure from intercellular airspace, C_i_ to that of the mesophyll, C_m_ are related in the following equation

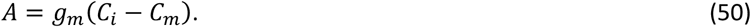

Incorporating equation (50) into equation (21) results in a cubic expression which is not easily solved. It can be incorporated into equation (40) giving a slightly more complex quadratic. In case of the initial slope of the CO_2_ response curve one can use equation (26) and ignoring the term g_s_C_m_ and combining it with equation (50) a quadratic similar to the one given for C_3_ leaves is obtained (von Caemmerer, 2000; von Caemmerer and Evans, 1991).

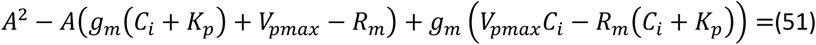

The first derivative with respect to C_i_ at C_i_=0 is given by

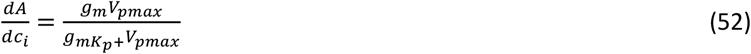

Pfeffer and Peisker (1988 and 1998) used this equation together with measurements of PEP carboxylase activity and initial slope (dA/dC_i_) to estimate g_m_ in plants grown under different light intensities. Figure 4 shows the effect inclusion of mesophyll conductance has on the initial slope. Equation 52 was used by Ubierna et al. (2016) to estimate g_m_ from in vitro measurements of PEP carboxylase activity, V_pmax,_ and they found good agreement with estimates of g_m_ from measurements of C^18^O^16^O isotope discrimination, however the uncertainties surrounding estimates of K_p_ discussed above need to be considered.

Strong correlations between leaf nitrogen, CO_2_ assimilation rate and PEP carboxylase activity have been observed in several studies (Meinzer and Zhu, 1998; Sage and Pearcy, 1987; Usuda, 1984; Wong *et al*., 1985) however it has been more difficult to provide quantitative correlations. Without the inclusion of a mesophyll conductance estimates of V_pmax_ form the initial slope are often less than what is measured in vitro. For example if the initial slope of the lower curve in Figure 4a is used to estimate V_pmax_ with an infinitely large g_m_ the predicted V_pmax_ is 58 µmol m^−2^ s^−1^ whereas it has here been modelled with a V_pmax_ of 200 µmol m^−2^ s^−1^ and gm = 1 mol m^−2^ s^−1^ bar^−1^. Hence mesophyll conductance is an important parameter in linking C_4_ biochemistry with gas exchange.

### Modelled light response of CO_2_ assimilation

It is well recognised that the light response of C_4_ photosynthesis does frequently not saturate (Cousins *et al*., 2006; Leakey *et al*., 2006). Figure 5 shows typical modelled light response curves of CO_2_ assimilation rate at several mesophyll CO_2_ partial pressures. In the current parametrisation CO_2_ assimilation rate is electron transport limited at all irradiances above C_m_=150 µbar at 25 °C. The shapes of the curves are determined by equation (34) which as for the C_3_ photosynthetic model remains empirical and the partition partitioning of electron transport between C_4_ and C_3_ cycle has been set at x=0.4 (equation 29). Furbank and von Caemmerer gave a detailed discussion about the optimal partitioning of electron transport capacity between C_3_ and C_4_ cycle (Peisker, 1988; von Caemmerer, 2000; von Caemmerer and Furbank, 1999). It is noteworthy that the fraction of electron transport allocated to the C_4_ cycle, x, equals 0.4 over a wide range of irradiances but drops at very low irradiance. Under low light the bundle-sheath CO_2_ partial pressures are close to the mesophyll CO_2_ partial pressure and electron transport is required for recycling of photo respiratory CO_2_. The optimal partitioning increases from 0.404 to 0.417 if oxygen is evolved in the bundle sheath (α=1). It also declines slightly with increasing temperature as Rubisco specificity for CO_2_ decreases (Jordan and Ogren, 1984; Sharwood *et al*., 2016).

**Figure 5.**
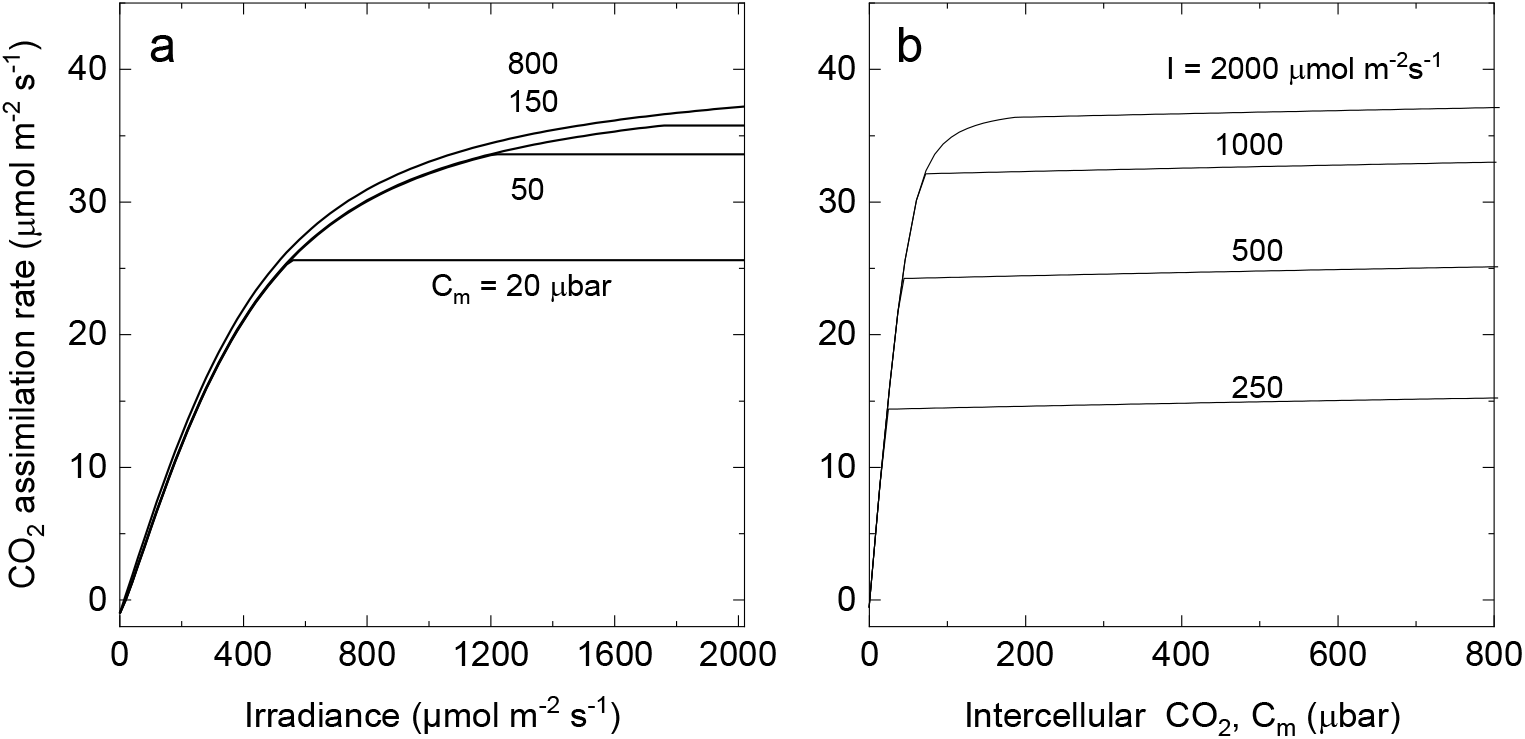
The effect of irradiance on modelled rate of CO_2_ assimilation. c) Light response of CO_2_ assimilation rate at mesophyll cytosolic CO_2_ partial pressures, C_m_, in dictated in the figure. The C_4_ photosynthesis model predicts electron transport limitations at all irradiance at C_m_ values above 150 µbar. At lower C_m_, CO_2_ assimilation rates are enzyme limited at high irradiance. The model was parameterised at 25 °C with values given in Table 1 and x=0.4. d) CO_2_ assimilation rate as a function of intercellular CO_2_ at irradiances indicated. The model was parameterised at 25 °C with values given in Table 1 and x=0.4.

In C_3_ species there a close link has been established between chloroplast electron transport capacity and electron transport chain intermediates such as cytochrome f (Yamori *et al*., 2010). In C_4_ photosynthesis this quantitative link between cytochrome f content and electron transport capacity needs also to be investigated.

### Modelled temperature response of CO2 assimilation rate

C_4_ plants have higher CO_2_ assimilation rates at high temperatures and higher photosynthetic temperature optima than their C_3_ counterparts largely because of the elimination of photorespiratory CO_2_ losses. (Berry and Björkman, 1980; Long, 1999). The temperature response of electron transport is not well characterised in C_4_ species. With the parameterisation used here, CO_2_ assimilation rate is electron transport limited above 25°C, and enzyme limited below a Ci=150 µbar at hight light, which corresponds to the operating C_i_ of many C_4_ species at ambient CO_2_ (Figure 6). There is some evidence that this is not unreasonable. Figure 7 shows a comparison of a temperature response of CO_2_ assimilation rate of *Flaveria bidentis* wildtype and transgenic Flaveria with reduced Rubisco content (Kubien *et al*., 2003). In Figure 7b CO_2_ assimilation rate is expressed on a Rubisco site basis (in vivo k_cat_) and compared the temperature response of Rubisco in vitro activity. For Flaveria with reduced amount of Rubisco there is a match between in vivo and in vitro k_cat_ up to approximately 30°C whereas for the wild type in vivo k_cat_ is less than the in vitro Rubisco k_cat_ around 20°C indicating other limitations to CO_2_ assimilation rate such as electron transport capacity. Temperature optima of CO_2_ assimilation rate are dependent on growth environment (Berry and Björkman, 1980; Dwyer *et al*., 2007). The modelling suggests that the temperature optimum is most likely determined by the properties of the electron transport capacity (Figure 6).

**Figure 6.**
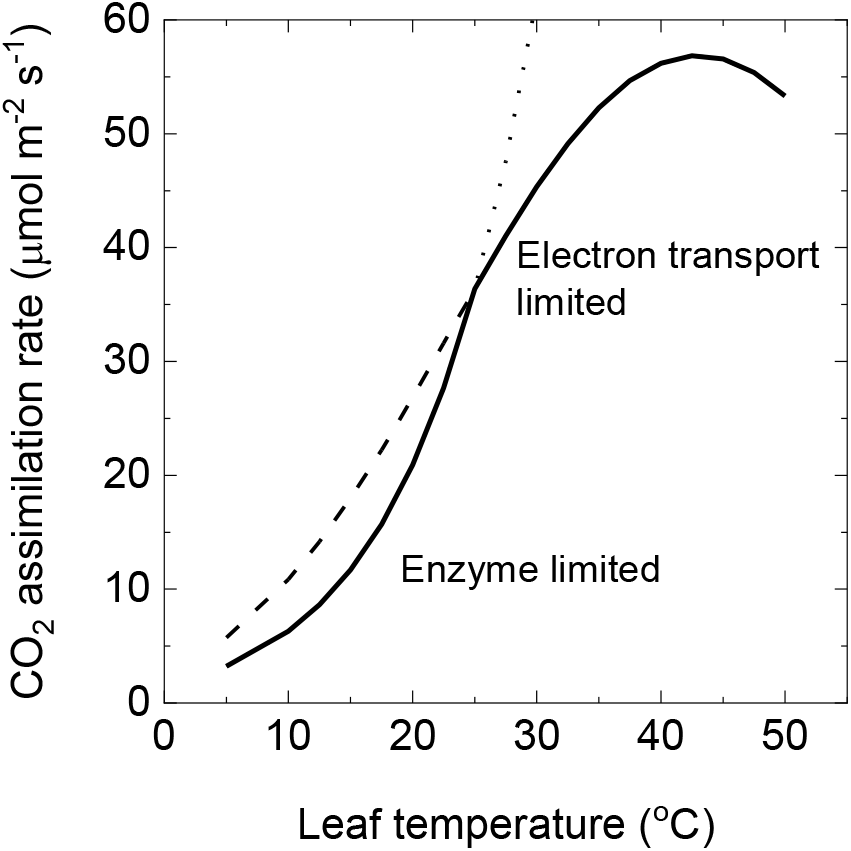
Modelled CO_2_ assimilation rate as a function of leaf temperature. The dotted line and its extension line show the enzyme limited rate and the dashed line and its extension line show the electron transport limited rate. CO_2_ assimilation rate was modelled at an irradiance of 2000 µmol m^−2^ s^−1^ and mesophyll CO_2_, C_m_ of 150 µbar. Other parameters are as given in Table 1.

**Figure 7.**
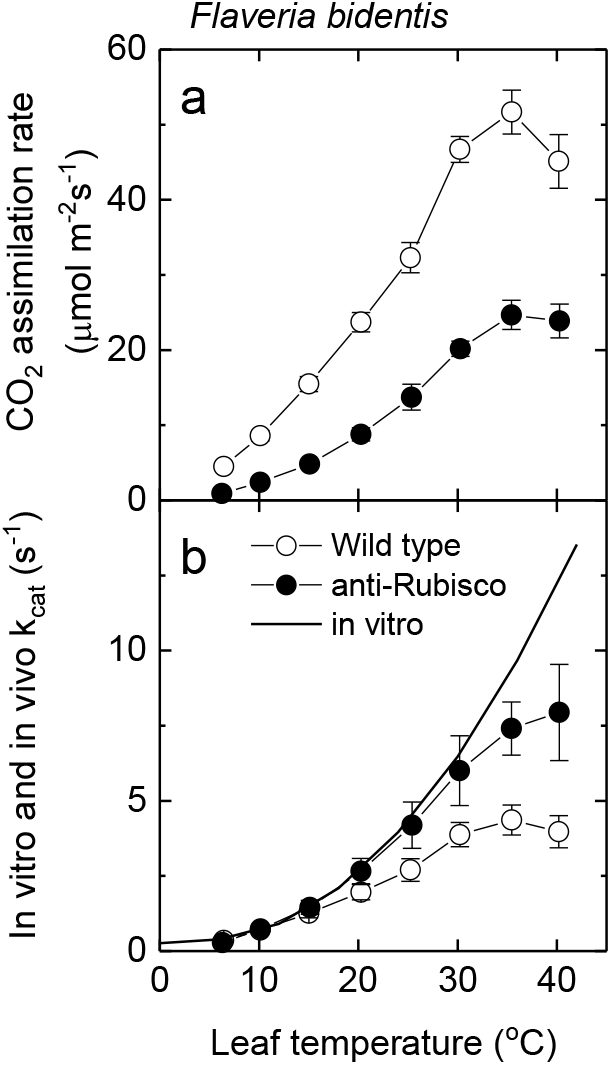
a) Temperature responses CO_2_ assimilation rate in *Faveria bidentis* wild type and anti-Rubisco plants Photosynthesis was measured at different leaf temperatures and ambient CO_2_ of 370 µbar and 200 mbar O_2_ and an irradiance of 1500 µmol m^−2^ s^−1^. Each point represents the mean (± SE) of measurements on five different leaves. b) Temperature dependence of the in vitro and in vivo k_cat_ for Rubisco in wildtype and anti-Rubisco *F. bidentis*. The in vitro data reflect the activity of the fully carbamylated enzyme; in vivo k_cat_ is estimated as gross photosynthesis divided by the number of Rubisco catalytic sites. Each value represents the mean (± SE) of four measurements. The data is redrawn from Figures 1 and 4 (Kubien *et al*., 2003)

In Figure 8a the CO_2_ response curves have been modelled for different leaf temperatures. Figure 8a, shows that at low temperature the CO_2_ response is enzyme limited at all Ci, as temperature is increased the transition from enzyme limited CO_2_ assimilation rate to electron transport limited rate occurs at progressively lower C_i_. There is an increase in initial slope with increasing temperature which is caused by the temperature response of the mesophyll conductance and the different temperature dependencies of maximal PEPC carboxylation and is Michaelis Menten constant (V_pmax_ and K_p_) (Figure 8 a and b, Table 1). These model predictions fit well with experimental observations by Sonawane et al.(2017).

**Figure 8.**
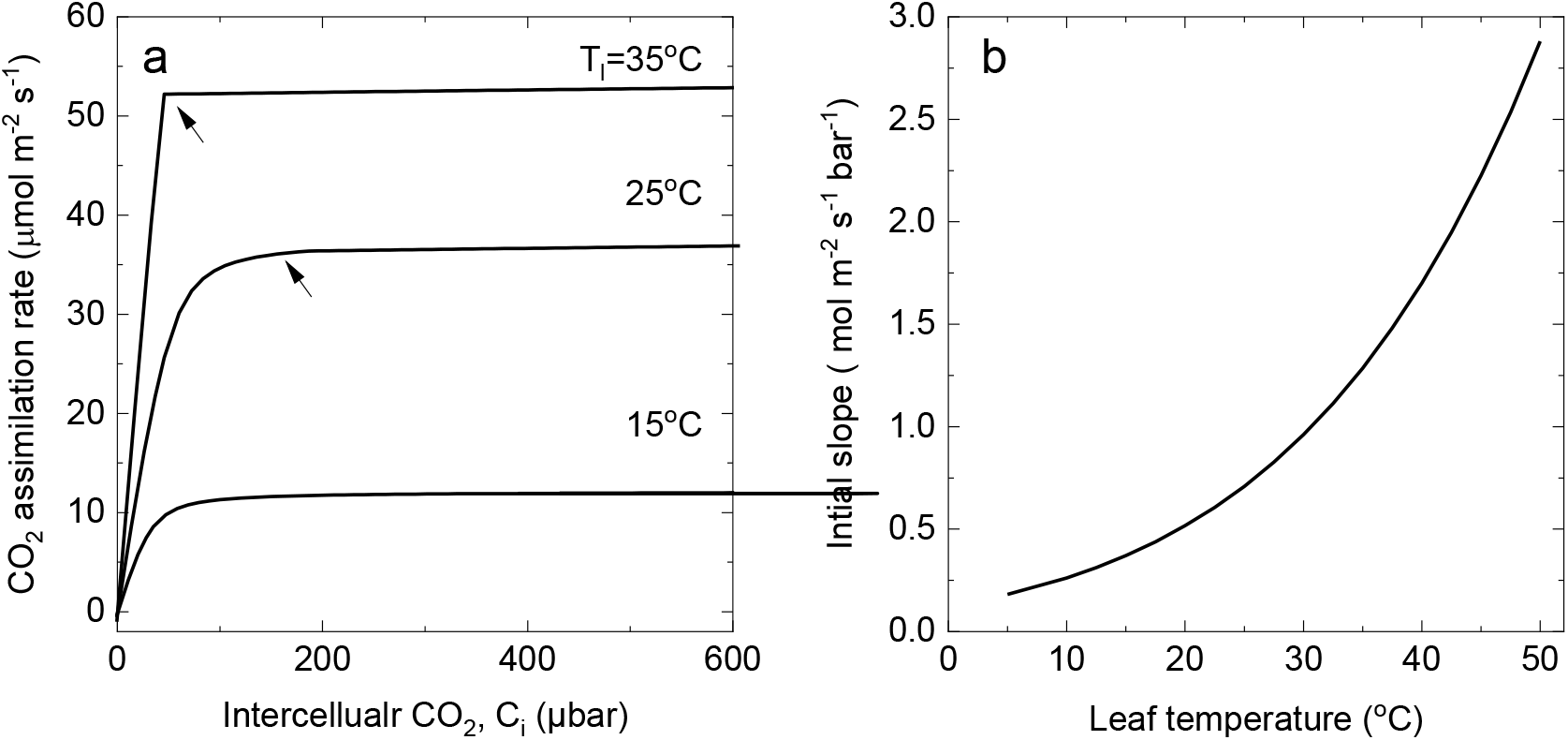
a) Modelled rate of CO_2_ assimilation as a function of intercellular CO_2_ partial pressure, C_i_, for the C_4_photosynthetic pathway at three leaf temperatures of 15, 25 and 35°C and an irradiance of 2000 µmol m^−2^ s^−1^. At 15°C CO_2_ assimilation rate is enzyme limited at all C_i_. For 25 and 35°C the arrow indicates the transition from enzyme limitation at low C_i_ to electron transport limitation at high C_i_. Parameters used are given in Table 1. b) Intial slope (dA/dCi) calculated from equation 52 as a function of leaf temperature. Parameters used are given in Table 1.

### A note on leakiness

The bundle sheath resistance or its inverse the bundle sheath conductance to CO_2_ diffusion are key parameters that together with relative capacities for the C_4_ cycle and Rubisco and electron transport capacity determine the effectiveness of the CO_2_ concentration mechanism. This is often quantified by a term called leakiness (*ϕ*), which is defined as the ratio of the rates of CO_2_ leakage out of the bundle sheath over the rate of CO_2_ supply to the bundle sheath (equation 4 and 5). Carbon isotope discrimination can be used to determine leakiness (Farquhar, 1983). Combined measurements of gas exchange and carbon isotope discrimination have been used to assess leakiness under different environmental conditions (Henderson *et al*., 1992; King *et al*., 2012; Kromdijk *et al*., 2010; Pengelly *et al*., 2010; Sun *et al*., 2012; Ubierna *et al*., 2011; von Caemmerer and Furbank, 2003). It is tempting to predict leakiness from the C_4_ photosynthesis model, but because it is a flux model little can be said about the rate of the component that is not limiting and leakiness estimates are not realistic as non-rate limiting steps are likely to be down regulated. However, when CO_2_ assimilation rates and leakiness are known from combined measurements of gas exchange and carbon isotope discrimination the model can be used to calculated the rate of the C_4_ cycle and the leak rate from the following equations and equation 4

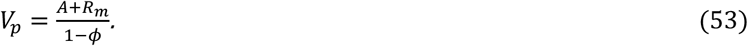

With the assumption of a bundle sheath conductance bundle sheath CO_2_ can also be estimated (Pengelly *et al*., 2012).

## Conclusion

The steady state C_4_ photosynthesis model has been updated and parameterized with the *in vitro* kinetic constants for Rubisco and PEP carboxylase and values for mesophyll and bundle sheath conductance and their temperature dependencies. Furthermore, electron transport rate equations have been updated to include cyclic electron transport flow. Now it is important to compare gas exchange measurements and biochemical measurements to confirm the quantitative relationships predicted by the model. In particular a parameterization of the temperature response of electron transport rate is needed and information of how it relates to thylakoid electron transport components such as the b_6_f complex which has shown to be a good correlator of C_3_ photosynthetic electron transport.

## Acknowledgment

I thank Robert T Furbank and Alex Wu for helpful comments on this manuscript. This research was supported by the Australian Research Council (ARC) Centre of Excellence for Translational Photosynthesis (CE140100015).

